# The node of Ranvier influences the *in vivo* axonal transport of mitochondria and signalling endosomes

**DOI:** 10.1101/2024.06.04.597271

**Authors:** Andrew P. Tosolini, Federico Abatecola, Samuele Negro, James N. Sleigh, Giampietro Schiavo

## Abstract

Efficient long-range axonal transport is essential for maintaining neuronal function, and perturbations in this process underlie severe neurological diseases. We have previously demonstrated that signalling endosomes are transported *in vivo* at comparable speeds across motor neurons (MNs) innervating different hindlimb muscles, as well as between forelimb and hindlimb peripheral nerves. In contrast, axonal transport is faster in MNs compared to sensory neurons innervating the same muscle. Found periodically across the myelin sheath, Nodes of Ranvier (NoR) are short uncovered axonal domains that facilitate action potential propagation. Currently, it remains unresolved how the distinct molecular structures of the NoR impact axonal transport dynamics. Here, using intravital time-lapse microscopy of sciatic nerves in live, anaesthetised mice, we assessed diverse organelle dynamics at the NoR. We first observed that axonal morphologies were similar between fast and slow MNs, and found that signalling endosomes and mitochondria accumulate on the distal side of the NoR in both motor neuron subtypes. Assessment of axonal transport of signalling endosomes and mitochondria revealed a decrease in velocity and increase in pausing as the organelles transit through the NoR, followed by an increase in speed in the adjacent intranodal region. Collectively, this study has established axonal transport dynamics of two independent organelles at the NoR *in vivo*, and has relevance for several pathologies affecting peripheral nerves and the NoR, such as peripheral neuropathy, motor neuron diseases, and/or multiple sclerosis.

## Introduction

Axonal transport is a fundamental biological process that maintains neuronal homeostasis with constant bidirectional shuttling of essential cargoes and structural components between the neuronal cell body and axon termini. Along the microtubule network, cytoskeletal elements (e.g., neurofilaments), as well as membranous (e.g., mitochondria, endosomes) and membrane-less (e.g., RNA granules) organelles undergo plus-ended directed anterograde transport driven by members of the kinesin motor protein family, and minus-end directed retrograde transport via cytoplasmic dynein (Maday et al., 2014; Vargas et al., 2022). Effective long-range axonal transport is essential for maintaining neuronal function, and trafficking perturbations underlie several neurodevelopmental and neurological conditions (Sleigh et al., 2019).

α-motor neurons (MNs) can be subclassified into fast MNs (FMNs) and slow MNs (SMNs), that selectively innervate specific skeletal muscle fibre subgroups. Both types of MN and the muscle fibres they innervate have distinct metabolic, functional and transcriptional properties (Stifani, 2014; Blum et al., 2021). Compared to SMNs that innervate slow oxidative type I muscle fibres, FMNs have larger motor unit sizes, faster nerve conduction, different firing patterns, and they innervate type IIa fast oxidative-glycolytic, as well as type IIx and IIb fast glycolytic muscle fibres (Stifani, 2014; Ragagnin et al., 2019).

We have previously determined that signalling endosome transport speeds are similar between motor axons innervating prototypical fast and slow muscles (Tosolini et al., 2022). Likewise, axonal transport speeds are comparable between hindlimb and forelimb peripheral nerves (Lang et al., 2023). In contrast, signalling endosome transport in MNs is faster than in sensory neurons innervating the same muscle (Sleigh et al., 2020a). Several factors directly influence axonal transport dynamics (Maday et al., 2014; Gibbs et al., 2015; Nirschl et al., 2017; Cason and Holzbaur, 2022), and can be perturbed in disease (Sleigh et al., 2019; Berth and Lloyd, 2023). Collectively, this suggests that the axonal transport machinery is differentially modulated in physiological and pathological contexts.

As most *in vivo* axonal transport studies have prioritised larger and/or consistently sized axonal segments, our understanding of transport dynamics at highly specialised axonal regions, such as the node of Ranvier (NoR), remains incomplete. NoRs are short uncovered axonal domains that facilitate action potential propagation, and each of the four regions (i.e., node, paranode, juxtaparanode and internode) are comprised of distinct structural and functional proteins (D’Este et al., 2017; Rasband and Peles, 2021). Electron microscopy reveals reductions in axonal diameters at the NoR, which maximise electrical conduction velocities (Johnson et al., 2015); these constrictions were attributed to reduced neurofilament, but not microtubule, content (Reles and Friede, 1991; Berthold et al., 1993; Hsieh et al., 1994). Most internodal neurofilaments are not continuous through the NoR and cease near the juxtaparanode, whereas microtubules extend through the NoR to connect adjacent internodes (Tsukita and Ishikawa, 1981), providing structural continuity for the continuous processive movement of cargoes.

Despite high organelle content (Berthold et al., 1993; Fabricius et al., 1993), unique axonal Ca^2+^ dynamics (Zhang et al., 2010) and high metabolic needs (Chiu, 2011), which are all features with the potential to regulate axonal transport, few studies have investigated transport dynamics at the NoR. Neurofilaments undergoing slow axonal transport (Brown, 2003) increase their speed through the NoR (Walker et al., 2019; Ciocanel et al., 2020). In contrast, muscle-administered horseradish peroxidase (HRP) (Berthold and Mellström, 1982), radiolabelled glycoproteins (Armstrong et al., 1987), and lysosome-linked enzymatic activity (Gatzinsky and Berthold, 1990) are enriched at peripheral nerve NoRs, but little is known about their transport.

The aim of this study was to assess the *in vivo* axonal transport of two different organelles at the NoR in FMNs and SMNs to further our understanding of fast axonal transport dynamics through this specialised structure. Knowledge of how the node affects cargo delivery may reveal key mechanisms relevant for pathologies affecting peripheral nerves.

## Materials and Methods

### Animals

Animal experiments performed in the United Kingdom were conducted in accordance with the European Community Council Directive of 24 November 1986 (86/609/EEC), under license from the UK Home Office in accordance with the Animals (Scientific Procedures) Act 1986, and were approved by the UCL Institute of Neurology Ethical Review Committee. Procedures carried out in Italy were approved by the ethical committee and by the animal welfare coordinator of the OPBA from the University of Padua. All procedures are specified in the projects approved by the Italian Ministry of Health, Ufficio VI (authorisation numbers: 359/2015PR; 81/2017 PR; 521/2018 PR; 439/2019 PR) and were conducted in accordance with National laws and policies (D.L. n. 26, March 14, 2014), following the guidelines established by the European Community Council Directive (2010/63/EU) for the care and use of animals for scientific purposes.

3-6 month old heterozygous male and female mice of the following transgenic strains were used: 1) Tg(Chat-EGFP)GH293Gsat/Mmucd mice (RRID: MMRRC_000296-UCD; Gong et al., 2003), referred to as ‘ChAT.eGFP’ mice; 2) B6.Cg-Tg(Thy1-CFP/COX8A)S2Lich/J (RRID: IMSR_JAX: 007967) Misgeld et al., (2007), referred to as ‘Mito.CFP’ mice; and 3) B6;CBA-Tg(Plp1-EGFP)10Wmac/J (RRID: IMSR_JAX:033357) (Mallon et al., 2002), referred to as ‘PLP-GFP’ mice. Mice were housed in individually ventilated cages in a controlled temperature/humidity environment and maintained on a 12 h light/dark cycle with access to food and water *ad libitum*.

### Intramuscular injections of HcT

Fluorescently-labelled atoxic fragment of tetanus neurotoxin (HCT-555) was prepared as previously described (Restani et al., 2012). Briefly, HCT (residues 875–1315) fused to an improved cysteine-rich region was expressed in bacteria as a glutathione-S-transferase fusion protein, cleaved and subsequently labelled with AlexaFluor555 C2 maleimide (Thermo Fisher Scientific, A-20346). Mice were anaesthetised using isoflurane, and after the fur on the ventrolateral lower leg was shaved, mice were placed on a heat-pad ready for intramuscular injections. A small incision was made on the ventral surface below the patella for the tibialis anterior muscle (TA), whereas for the soleus muscle a vertical incision was made on the skin covering the lateral surface of lower hindlimb between the patella and tarsus to expose the underlying musculature. Guided by previously established motor end plate maps (Mohan et al., 2014), intramuscular injections were performed, with 7.5-10 μg of HCT in PBS in a volume of ∼3.5 µl using a 701 N Hamilton® syringe (Merck, 20,779) for TA, or 1 µl in PBS using pulled graduated, glass micropipettes (Drummond Scientific,5-000-1001-X10) for soleus, as previously described (Mohan et al., 2015). Upon HCT administration, the skin was sutured, and mice were monitored for up to 1 h before returning to the home cage.

### In vivo imaging

Signalling endosomes were visualised *in vivo* after administration of HCT, as previously described (Gibbs et al., 2016; Sleigh et al., 2020b; Tosolini et al., 2021). At least 4 h after HcT intramuscular injections, mice were re-anaesthetised with isoflurane, and the sciatic nerve was exposed by first removing the skin and then the overlying biceps femoris muscle. To aid the imaging process a small piece of parafilm was inserted between the underlying connective tissue and the sciatic nerve. The anaesthetised mouse and nosepiece were then transferred to an inverted LSM780 confocal microscope (Zeiss) enclosed within an environmental chamber maintained at 37°C. Time-lapse microscopy was performed using a 40×, 1.3 NA DIC Plan-Apochromat oil-immersion objective (Zeiss) focusing on axons around the NoR using an 80× digital zoom (1024 × 1024, < 1% laser power). Frame intervals of ∼0.4-0.5 s were used when acquiring transport videos of motile HCT-positive signalling endosomes, whereas frame intervals of ∼1.5-2.0 s were used when acquiring videos of motile Mito.CFP-positive mitochondria alone or in combination with HCT-positive signalling endosomes. All imaging was concluded within a maximum of 1 h from initiating re-anaesthesia.

### Morphological analysis of nodes of Ranvier

Axonal morphologies at the NoR were assessed using the HCT transport videos obtained in TA- or soleus-innervating motor axons from ChAT.eGFP mice, as previously described (Sleigh et al., 2020a; Tosolini et al., 2022). Orthogonal measurements were made between the upper and lower or proximal and distal ends of axonal regions containing motile HCT signalling endosomes (**Figure 1A**). A minimum of ten measurements (i.e., combined proximal and distal internodal axonal diameter, axon diameter in the nodal constriction, and axon length in the nodal constriction) from at least three different axons were used to calculate the average diameter/length per animal.

**Figure 1.**
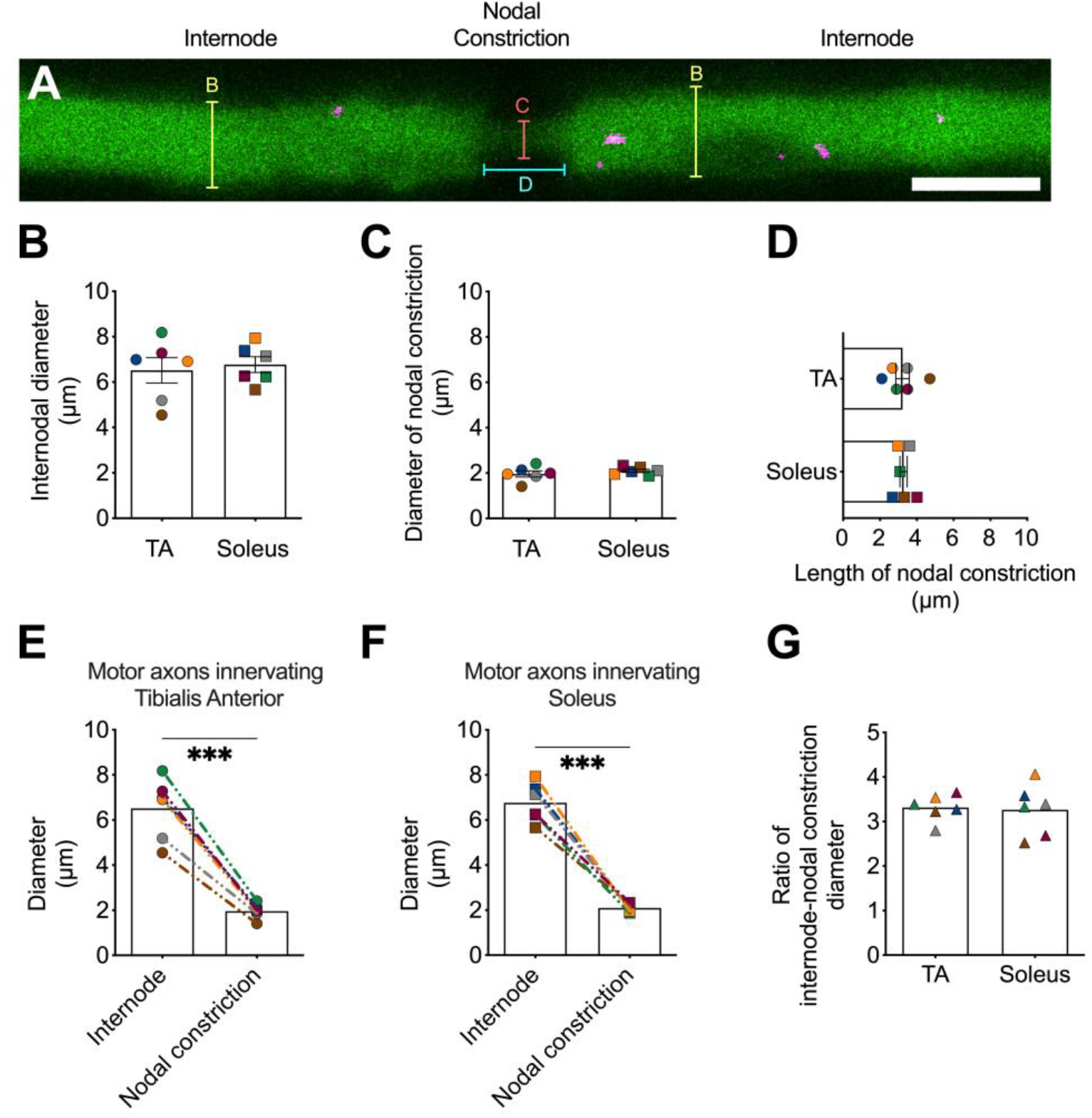
Morphology of the Node of Ranvier is similar between fast and slow motor axons of the sciatic nerve. **A)** Representative single frame image of HCT-555 containing signalling endosomes (magenta) in a single motor axon (green) from a ChAT.eGFP mouse sciatic nerve. Scale bar = 5 µm. Yellow lines represents the axonal region assessed to quantify the internodal diameters in **B**, the orange line represents the axonal region assessed to quantify the nodal constriction in **C**, and the cyan line represents the axonal region assessed to quantify the length of the nodal constrictions in **D**. Motor axons innervating tibialis anterior (TA) or soleus muscles display similar **B)** internodal diameters (p = 0.71, unpaired two-tailed t test), **C)** nodal constriction diameters (p = 0.40, unpaired two-tailed t test), and **D)** lengths of nodal constriction (p = 0.90, unpaired two-tailed t test). Furthermore, the differences in axonal diameters at the internode and nodal constriction were comparable between motor axons innervating **E**) TA and **F**) soleus muscles, and the **G)** ratios of internodal to nodal diameters were also comparable (p = 0.85, unpaired two-tailed t test). ***p <0.001 assessed by two-tailed paired t test. For all graphs, n=6, means ± SEM are plotted, and the colour-coding remains consistent between animals. **Linked to Movie 4**.

### Analysis of the distribution of signalling endosomes and mitochondria

The distributions of signalling endosomes and mitochondria at the NoR were assessed from the axonal HCT transport videos in TA- or soleus-innervating motor axons in Mito.CFP mice. Using FIJI/ImageJ, the relative fluorescence profiles of HCT-containing signalling endosomes and mitochondria across the NoR and internodal regions were measured after drawing an 80 µm region centred around the nodal constriction from the Z-stack projection. After the background was subtracted, the relative fluorescence intensity profiles were then plotted, and averaged from a minimum of 3 axons per animal.

### Immunohistochemistry of the sciatic nerve

#### NoR staining

Sciatic nerves were dissected from euthanised ChAT.eGFP mice and fixed in 4% paraformaldehyde in PBS for approximately 1 h at room temperature. Sciatic nerves were washed in PBS three times for 5 min, teased into individual fibres/bundles and were then permeabilised and blocked in a blocking solution containing 1% Triton X-100 and 10% bovine serum albumin in PBS for 1 h at room temperature. Sciatic nerve fibres were then incubated in a blocking solution containing primary antibodies against NaV1.6 [SCN8a] (1:400; Alomone, ASC-009) and Caspr [clone K65/35] (1:500; Neuromab 75-001, UC Davies) for ∼3 d at 4°C with mild agitation. Following three washes in PBS at room temperature, fibres were then immersed in a solution containing anti-mouse-555 (1:500; Thermo Fisher Scientific; A-21422) and anti-rabbit-647 (1:500; Thermo Fisher Scientific; A31573) secondary antibodies in PBS for ∼2 h at room temperature. Fibres were then washed three times in PBS, mounted in Fluoromount-G (Thermo Fisher Scientific, 00-4958-02) and covered with 22 × 50 mm cover glass (VWR, 631-0137). Slides were dried and imaged with a LSM780 confocal microscope using a 63× Plan-Apochromat oil immersion objective (Zeiss).

#### HCT and Schwann cell cytosol

Anaesthetised PLP-GFP mice were injected into the TA muscle with HCT (Tosolini et al., 2021); after 24 h, mice were euthanised and the sciatic nerves dissected and fixed in 4% paraformaldehyde in PBS for 2 h at room temperature. After 3 x 5 min washes in PBS, sciatic nerves were then de-sheathed, teased into individual fibres and mounted using Dako fluorescence mounting medium (Agilent Technologies, S3023). Z-stack images were obtained with a confocal microscope (Zeiss LSM900 Airyscan2) equipped with an EC Plan-Neofluar 40×/1.30 oil objective.

### Tracking analysis

Confocal “.czi” images were opened in FIJI/ImageJ (http://rsb.info.nih.gov/ij/), converted to “.tiff”, and using the TrackMate plugin (Ershov et al., 2022) were tracked in a semi-automatic way for HcT-signalling endosomes (Tosolini et al., 2022) or manually for mitochondria (Kalinski et al., 2019). The following criteria were used for the tracking analysis: 1) only endosomes and mitochondria that were moving for ≥10 consecutive frames were included, and terminal pauses, as defined by the absence of movement in ≥ 10 consecutive frames, were excluded; 2) each axon required a minimum of 20 trackable organelles per axon, except for retrograde moving mitochondria which only required a minimum of 10 per axon; 3) at least three separate axons were assessed per mouse. A pause was defined by a previously motile organelle with a velocity of ≤ 0.1 µm/s between consecutive frames (to account for a potential breathing/arterial pulsing artefact), and the time paused (%) was determined by dividing the number of pauses by the total number of frame-to-frame movements assessed from an individual axon.

Transport dynamics of the retrogradely moving HCT signalling endosomes, as well as anterogradely and retrogradely moving mitochondria were separately assessed with TrackMate (Ershov et al., 2022), using the abovementioned criteria. Each frame-to-frame recording (e.g., velocity, pausing) was matched with its x and y co-ordinates, binned every 2 µm across an 80 µm axonal length centred at the NoR, and subsequently separated into either the proximal internode, nodal constriction, or distal internode locations. The mean moving velocity represents the average of all frame-to-frame movements in a particular axonal area (excluding pausing events), whereas the relative frequency of mean pausing was determined by assessing the relative number of pauses in relation to the axonal location.

### Statistics

GraphPad Prism 10 Software (Version 10.2.3) was used for all statistical analyses and figure production. Data were assumed to be normally distributed, and parametric data were assessed using paired or unpaired two-tail *t*-tests, as well as one-way or two-way analyses of variance (ANOVA) with Holm-Šídák’s multiple comparison tests.

## Results

### Identifying the node of Ranvier *in vivo*

To confirm the identity of the NoR, we performed immunohistochemistry on fixed, teased sciatic nerve fibres from ChAT.eGFP mice probing for established markers of the node and paranodal regions (Arancibia-Cárcamo et al., 2017). As expected, we observed a ring-shaped cluster of NaV1.6 channels approximately in the middle of the ChAT.eGFP nodal constriction that is flanked by CASPR immunolabelling of the paranode (**Movie 1**). Therefore, we concluded that the identical constrictions observed in ChAT.eGFP sciatic nerves contain the nodal and paranodal regions and can be distinguished from the larger internodal regions using intravital imaging of the endogenous eGFP fluorescence in ChAT.eGFP mice.

To locate the NoR in FMN and SMN axons *in vivo*, we performed intramuscular injections of HCT separately into TA or soleus muscles in ChAT.eGFP mice. HCT is internalised into axon termini of motor neurons through the binding to polysialogangliosides, nidogens (Bercsenyi et al., 2014) and the tyrosine phosphatases LAR and PTPRδ (Surana et al., 2024). Following sorting into a Rab5-positive endosomal compartment, HCT undergoes fast axonal retrograde transport toward the cell bodies of spinal cord motor neurons in Rab7-positive organelles (Deinhardt et al., 2006; Goto-Silva et al., 2019) using a cytoplasmic dynein- and microtubule-dependent process (Surana et al., 2018).

Intravital imaging of ChAT.eGFP sciatic nerve cholinergic axons (i.e., MN axons) enabled the identification of nodal constrictions that are bordered on both sides by wider internodal regions (**Fig. 1A**). Assessing individual axons from the TA-innervated FMNs, or soleus-innervated SMNs revealed similar diameters of the internodal regions (**Fig. 1B**) and nodal constrictions (**Fig. 1C**), as well as the lengths of the nodal constrictions (**Fig. 1D**). This analysis also revealed that the differences in axonal diameters at the internode and nodal constriction were comparable between motor axons innervating TA (**Fig. 1E**) and soleus (**Fig. 1F**), which was confirmed when the ratios of internodal to nodal diameters ratios were calculated (**Fig. 1G**). From these analyses, we concluded that the morphological features of the NoR are equivalent in wild-type peripheral nerve motor axons innervating prototypical fast and slow muscles.

#### Signalling endosomes and mitochondria cluster at the distal NoR

We next profiled the spatial location of signalling endosomes and mitochondria in relation to the nodal constriction in fast and slow motor axons. To label the different motor axon subtypes, we injected HCT into either TA or soleus muscles of Mito.CFP mice, and performed simultaneous time-lapse microscopy of HCT-signalling endosomes and mitochondria (Tosolini et al., 2021), focusing specifically on the NoR (**Fig. 2Ai**; **Movie 2**). The individual fluorescence intensity profiles of both organelles were first created using z-projections of individual frames of time-lapse videos (**Fig. 2Aii-iii)**, and then the fluorescence intensities of HCT-signalling endosomes and mitochondria were assessed across the whole region (80 µm). Surprisingly, we detected an accumulation of the fluorescent signal of both organelles specifically at the distal side of the nodal region (**Fig. 2B**). To quantify this, we directly compared the average axonal fluorescence intensities of signalling endosomes and mitochondria from individual videos of TA- and soleus-innervating axons from proximal and distal internodal regions. These analyses revealed that there were a greater number of immobile HCT-containing signalling endosomes (**Fig. 2C**) and mitochondria (**Fig. 2D**) at the distal side of the NoR, and this distribution was similar in TA- and soleus-innervating motor axons (**Fig. 2C,D)**. To rule out that this clustering phenotype was caused by the administration of HCT and generation of HCT-containing signalling endosomes *in situ*, we evaluated the mitochondrial fluorescence profiles in naïve Mito.CFP axons (i.e., without intramuscular HCT injections). This analysis revealed the same mitochondrial distribution as shown in **Fig 2D**, demonstrating that the accumulation of mitochondria seen distally at the NoR is independent of the presence of HCT (**Supplementary Figure 1)**.

**Figure 2.**
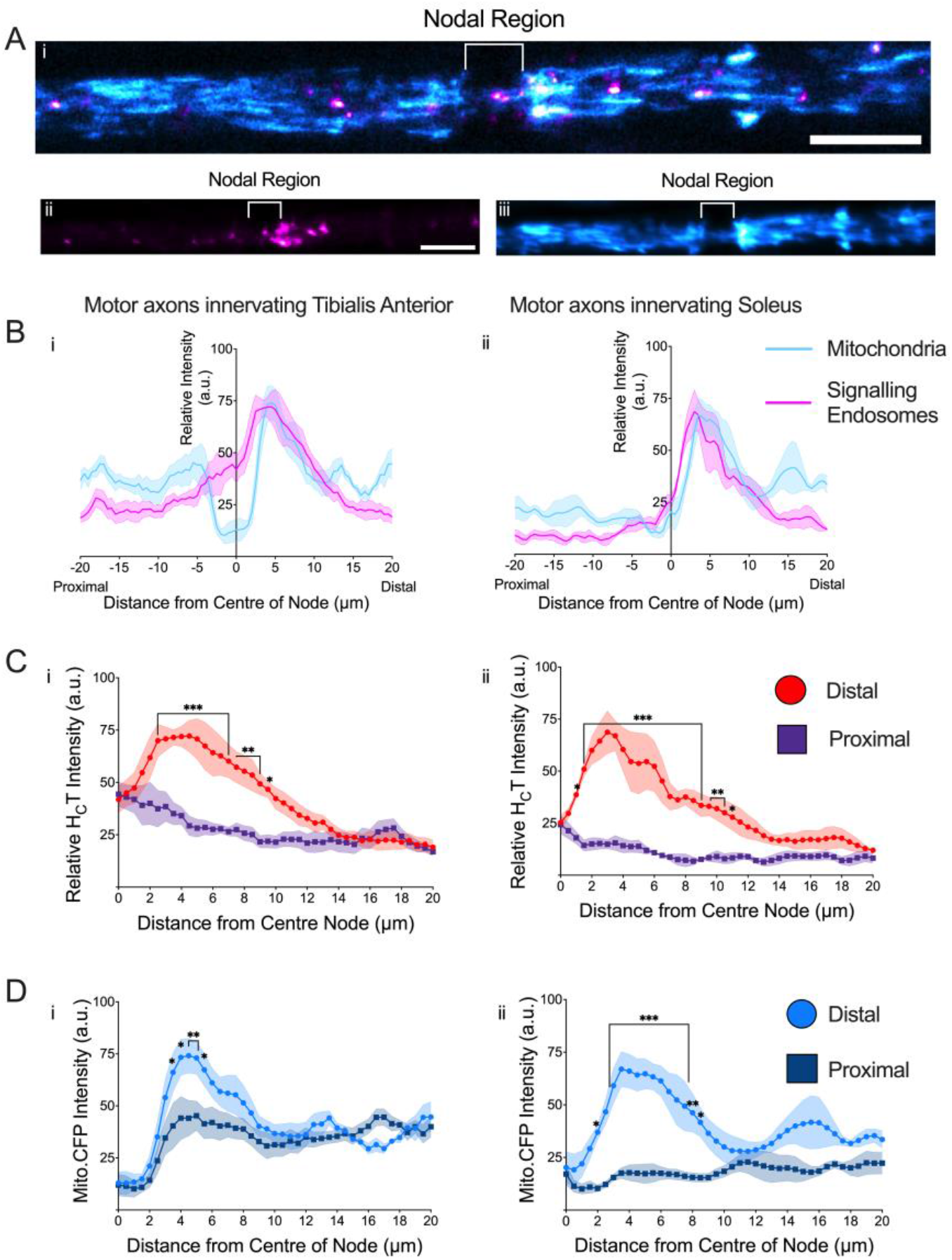
Signalling endosomes and mitochondria selectively accumulate at sites distal to the node of Ranvier. **A)** *i*. Representative image demonstrating the nodal distribution of HCT-containing signalling endosomes (magenta) and mitochondria (cyan) in a single tibialis anterior-innervating motor axon in a Mito.CFP mouse. Representative z-stack projection (sum of the slices) from a time-lapse video (see **Movie 2**) indicates that *ii)* HCT-containing signalling endosomes and *iii)* Mito.CFP labelled mitochondria are enriched on the distal side of the NoR. Scale bar = 10 μm. **B)** In motor axons innervating the *i*) Tibialis Anterior and *ii*) Soleus muscles, we observed increased average relative fluorescence intensities of both HCT-containing signalling endosomes (magenta) and Mito.CFP labelled mitochondria (cyan), exclusively on the distal side of the NoR. Enhanced fluorescence profiles of **C)** HCT containing signalling endosomes between 1-11 µm from the centre of the NoR (i. *TA* - Axonal Location: p <0.0001; Mean Relative Fluorescence: p <0.0001; Interaction: p <0.0001. ii. *Soleus:* Axonal Location: p <0.0001; Mean Relative Fluorescence: p <0.0001; Interaction: p <0.0001), and **D)** mitochondria 2-8.5 µm from the centre of the NoR (i. *TA* - Axonal Location: p <0.0001; Mean Relative Fluorescence: p <0.0001; Interaction: p = 0.0002. ii. *Soleus -* Axonal Location: p <0.0001; Mean Relative Fluorescence: p <0.0001; Interaction: p <0.0001). Means (solid line) ± SEM (shaded area) were plotted for all graphs, n=5 animals, 16 axons. Data were compared by two-way ANOVA and Holm-Šídák’s multiple comparisons tests. *p < 0.05, **p < 0.01, ***p < 0.001.

Previous studies suggest that axon-Schwann cell interactions regulate several critical functions. For example, CXCL12α/SDF-1 released from perisynaptic Schwann cells promotes motor axon regeneration (Negro et al., 2017), and ATP released from neurons activates Schwann cells (Rodella et al., 2017). We therefore aimed to evaluate whether HCT-positive signalling endosomes are released by exocytosis from this site of communication between motor axons and Schwann cells. To do so, we injected HCT into TA muscles in the PLP-GFP mouse, in which GFP is exclusively expressed in Schwann cells (Negro et al., 2022). 24 h after HCT injections, teased individual axons were isolated from the PLP-GFP sciatic nerves, and imaged. HCT clusters were again found in the distal portion of NoR; however no signalling endosomes were detected in the cytosol of Schwann cells (**Movie 3)**, ruling out that HCT transferred from motor axons to Schwann cells. Altogether, using three separate transgenic reporter mouse models, we have shown that HCT-positive signalling endosomes, along with mitochondria, cluster specifically at the distal side of the NoR.

#### Axonal transport of signalling endosomes and mitochondria through the node of Ranvier

We next sought to determine the axonal transport dynamics of both signalling endosomes and mitochondria through the NoR. First, we established that in ChAT.eGFP motor axons, HCT-containing signalling endosomes narrow their trajectories as they traverse the nodal constriction (**Fig. 3A; Movies 2 and 4**). We found that bidirectionally moving mitochondria also funnel their paths through the NoR when transitioning from the larger internodal regions through the nodal constriction, and then widen their trajectories when entering the subsequent internodal region (**Fig. 3B; Movie 2**). We also observed non-linear courses that individual HCT-positive signalling endosomes would take as they approached the nodal constriction (e.g., moving in an ‘S’ shape (**Fig. 4A; Movie 5)** or travelling in a circular motion (**Fig. 4B-C; Movie 6**)). These unusual transport trajectories might be caused by signalling endosomes traversing areas where the normal microtubule distribution observed in internodal regions is altered in proximity to nodal constrictions, e.g., radial axonal expansions (Wang et al., 2020), or circling areas with a high density of immobile organelles (e.g., mitochondria, **Fig. 4C**).

**Figure 3.**
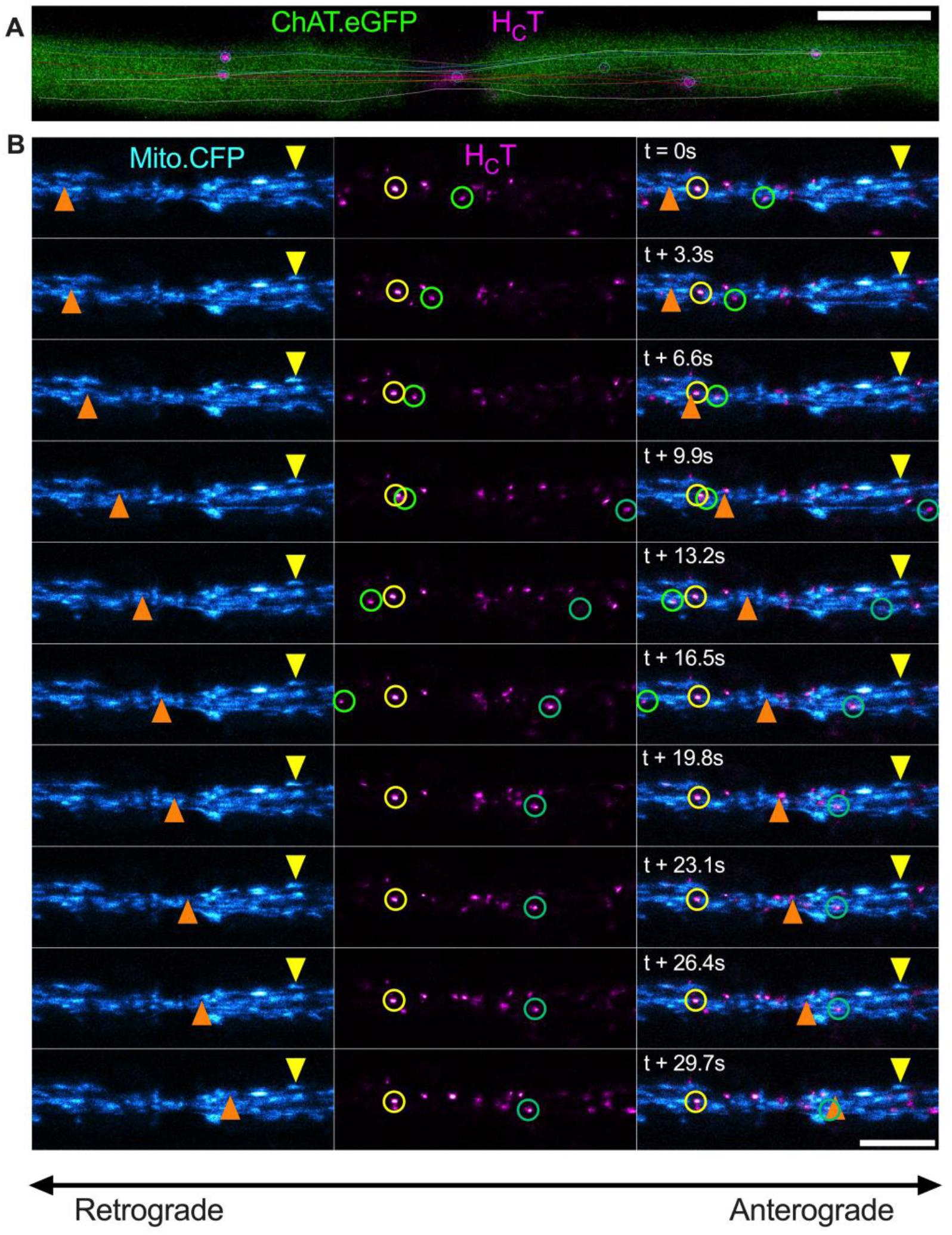
Axonal cargoes are bidirectionally funnelled through the node of Ranvier (NoR). **A)** *In vivo* imaging of the NoR reveals that trajectories (lines) of HCT-containing signalling endosomes (circles) narrow as they pass through the nodal constriction in sciatic nerve motor axons from a ChAT.eGFP mouse and expand in the adjacent internodal region. **B)** Time-lapse images taken every 3.3 s of mitochondria (cyan) and HCT-containing signalling endosomes (magenta) at the NoR from the same sciatic nerve axon in a Mito.CFP mouse. Turquoise and green circles identify retrogradely moving HcT-containing signalling endosomes, orange triangles identify anterograde moving mitochondria, and yellow triangles/circles identify paused/immobile organelles. Anterograde movement is from left to right, and retrograde movement is in the opposite direction. Scale bar = 5 μm.

**Figure 4.**
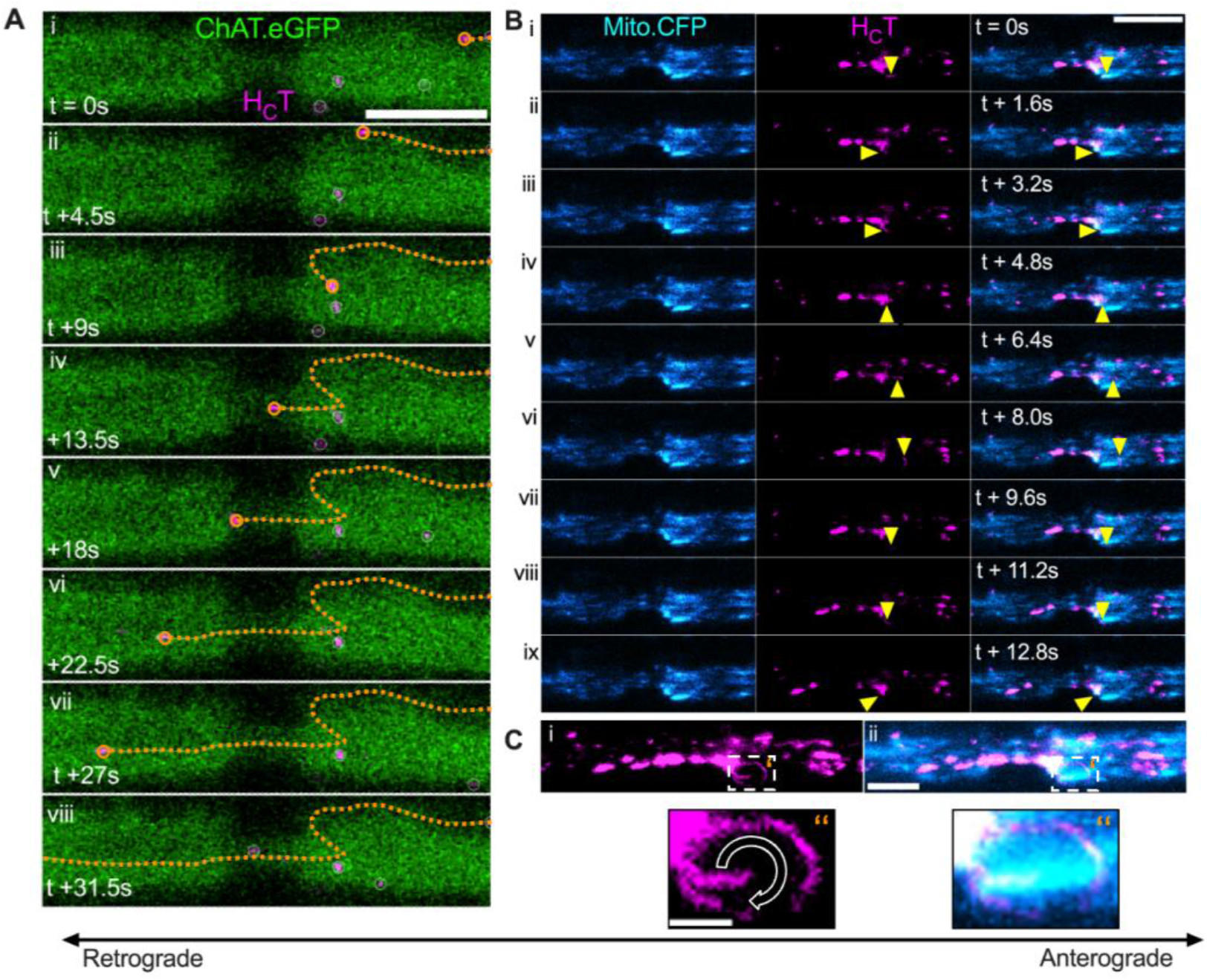
Non-linear trajectories of signalling endosomes are evident as they funnel through the node of Ranvier. **A)** Time lapse image series of an individual HCT-containing signalling endosome (magenta, with orange circle) travelling on an “S-shaped” trajectory (orange dotted lines) during *in vivo* axonal transport through the NoR from a ChAT.eGFP sciatic nerve motor axon. Frame interval = 4.5 sec. Linked with **Movie 5. B)** Time lapse image series demonstrating an individual HCT-containing signalling endosome (magenta, with yellow arrowheads) moving in a “circular” pathway around an individual mitochondrion (cyan) on the distal side of the NoR in a Mito.CFP sciatic nerve axon. Frame interval = 1.6 sec. Linked with **Movie 6. C)** Z-stack maximum projection of **B**, highlighting the circular pathway that an individual HCT-containing signalling endosome (magenta) as it traverses around a mitochondrion. C’’ are enlargements of the boxed area in C`’. Scale bar C = 10µm, C’’ = 2 μm.

Following these observations, we then used semi-automated tracking of HCT-signalling endosomes in TA- and soleus-innervating motor axons of the Mito.CFP mouse using the TrackMate plugin (Ershov et al., 2022) to assess transport dynamics. Strikingly, retrogradely moving HCT-containing signalling endosomes reduce their mean velocities when approaching the nodal constriction, with a concomitant increase in the relative frequency of pausing. This was followed by an increase in retrograde velocities on the proximal side of the NoR in both TA-(**Fig. 5A**) and soleus-innervating motor axons (**Supplementary Figure 2)**. Quantitative analyses in TA-innervating axons indicated that the mean velocities of HCT-containing signalling endosomes in the nodal constriction were ∼48% and ∼40% slower compared to the proximal and distal internodal regions, respectively (**Fig. 5B)**. Furthermore, the retrograde transport speeds through the proximal internode were ∼14% faster compared to distal internode regions (**Fig. 5B)**. In addition, in TA-innervating motor axons pausing was ∼86% and ∼67% lower in the proximal and distal internodal regions respectively, compared to the nodal constriction (**Fig. 5C**). Similarly in soleus-innervating motor axons, the mean velocities of HCT-signalling endosomes in the nodal constriction were ∼46% and ∼35% slower compared to the proximal and distal internodal regions, respectively (**Supplementary Figure 2B,C)**. As the transport dynamics of HCT-signalling endosomes at the NoR largely overlap in FMN and SMN motor axons (**Fig. 5D,E; Supplementary Figure 2 D,E)**, we conclude that motor neuron subtypes do not determine overt changes in axonal transport dynamics at the NoR, which is consistent with our previous observations across the larger internodal axonal region (Tosolini et al., 2022) To determine whether these phenotypes are either specific for the organelle and/or direction of travel, we assessed *in vivo* mitochondrial axonal transport in TA-innervating motor axons, from the same videos as the analysis of HCT-signalling endosomes (e.g., **Fig. 5**). Using the manual tracking feature of TrackMate (Ershov et al., 2022), we assessed axonal transport of mitochondria through the nodal constrictions separately for anterograde and retrograde directions (**Movie 7**). Like our observations for HCT-containing signalling endosomes, the mean velocity of mitochondria is also reduced at the nodal constriction, which was then followed by an increase in velocities for both anterograde and retrograde moving mitochondria (**Fig. 6A Supplementary Figure 3**). Anterograde mitochondrial velocities in the nodal constriction were ∼28% and ∼21% slower (**Fig. 6B**) compared to the proximal and distal internodes, respectively. Similarly, the mean velocities of retrogradely moving mitochondria were ∼24% and ∼23% slower in the nodal constriction compared to the proximal and distal internodal regions, respectively (**Fig. 6B**). We also observed faster mean velocity for anterograde moving mitochondria in the proximal internode compared to the distal internode, replicating the results obtained with signalling endosomes (**Fig. 5B**). However, unlike the significant increase in signalling endosome pausing frequency observed at the nodal constriction (**Fig. 5C)**, the increase in pausing frequency for anterograde and retrograde moving mitochondria is not statistically significant (**Fig. 6C)**.

**Figure 5.**
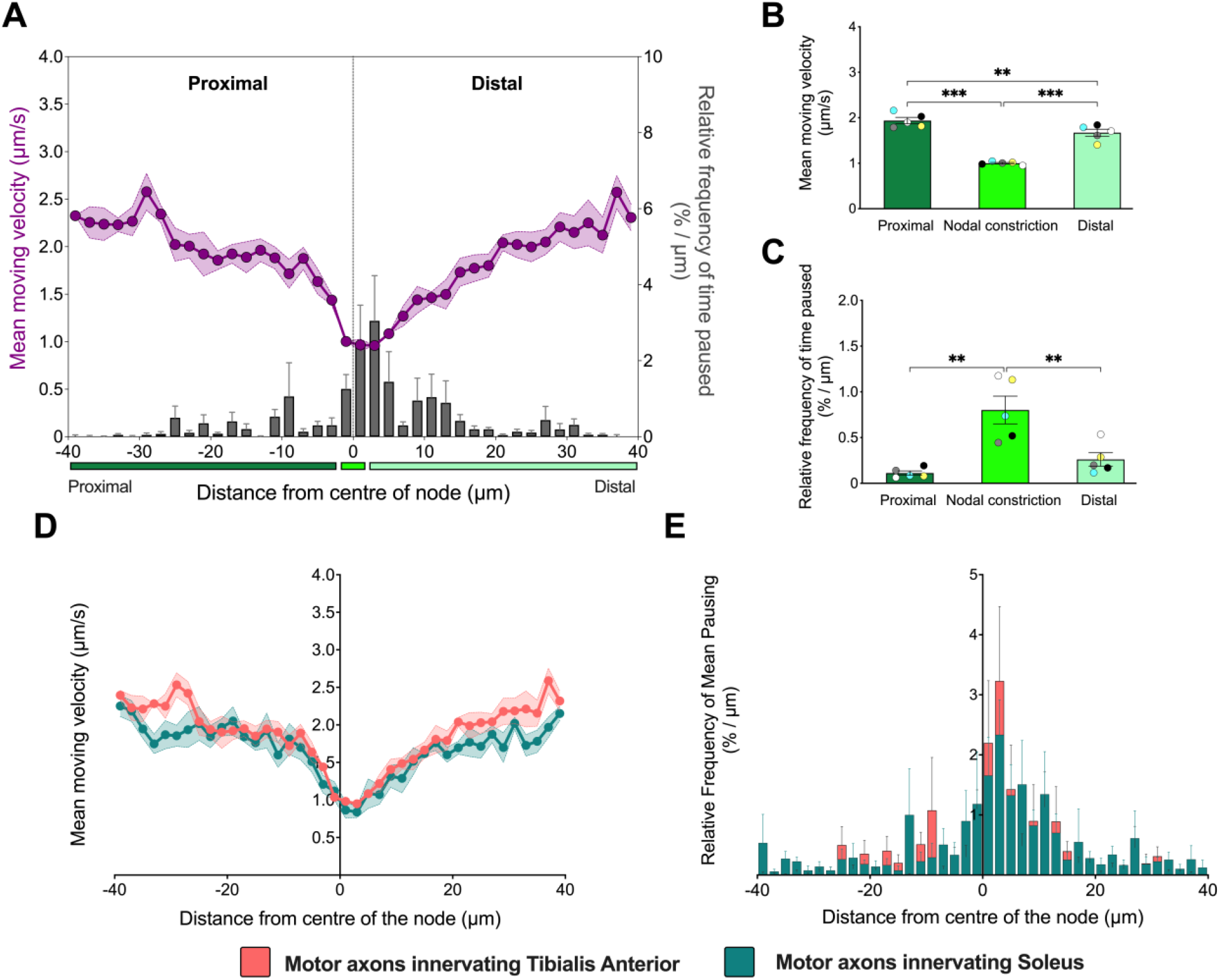
Signalling endosomes decelerate and pause more at nodes of Ranvier regardless of motor neuron subtype. **A)** Retrograde axonal transport dynamics (mean moving velocity [purple] and relative frequency of mean pausing [grey bars]) of HCT-containing signalling endosomes plotted across the nodal constriction (80 µm distance) in motor axons innervating the tibialis anterior muscle. X-axis represents the distance from the centre of the node of Ranvier (µm) and is split into three segments: 1) *Proximal* = 38 µm of the proximal internode (dark green); 2) *Centre* = 4 µm representing the mean nodal length, (as determined in **Figure 1D**; bright green); and 3) *Distal* = 38 µm of the distal internode (light green). Comparisons across the proximal internode (dark green), nodal constriction (bright green) and distal internode (light green) of the **B)** mean moving velocity (p < 0.001), and **C)** relative frequency of mean pausing (p = 0.0008). Data were compared by an ordinary one-way ANOVA, followed by Holm-Šídák’s multiple comparisons test. Histograms comparing the **D)** mean moving velocity and **E)** relative frequency of mean pausing from motor axons innervating the tibialis anterior (salmon) and soleus (teal). Means (solid line) ± SEM (shaded areas/error bars) were plotted for all graphs. n=4 (soleus), n=5 (tibialis anterior) from Mito.CFP mice.*p < 0.05, **p < 0.01, ***p < 0.001.

**Figure 6.**
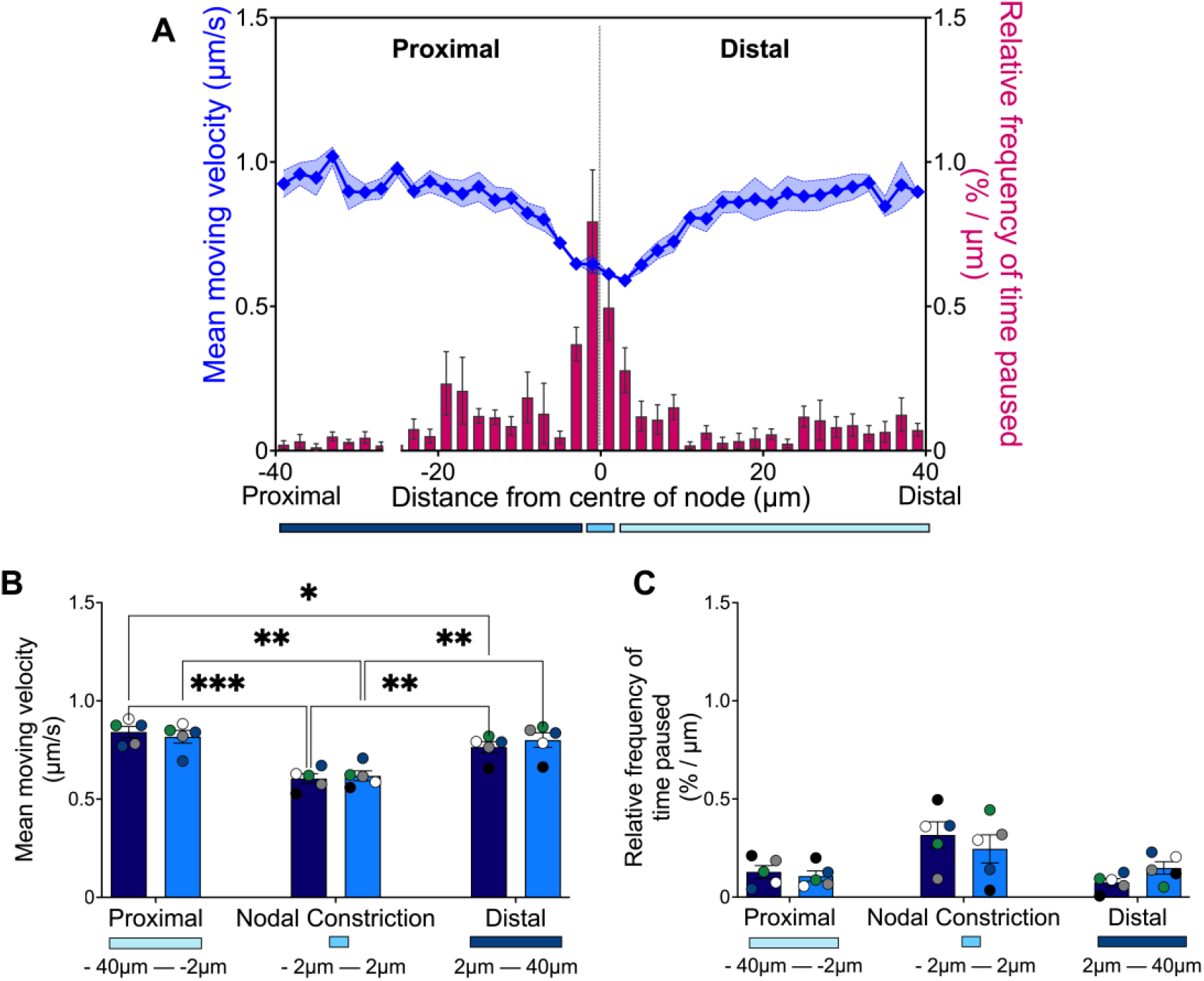
Mitochondria also decelerate and pause more at the node of Ranvier regardless of transport direction. **A)** Mitochondrial (i.e., anterograde + retrograde) axonal transport dynamics (mean moving velocity [blue] and relative frequency of mean pausing [pink bars]) across the nodal constriction (80µm distance) in sciatic nerve axons innervating the tibialis anterior muscle. X-axis represents the distance from the centre of the node of Ranvier (µm) and is split into three segments: 1) *Proximal* = 38 µm of the proximal internode (navy); 2) *Centre* = 4 µm representing the mean nodal length (as determined in **Figure 1D**; cyan); and 3) *Distal* = 38 µm of the distal internode (light blue). Comparisons between the nodal constriction, and proximal and distal internodes of *in vivo* axonal transport of anterograde and retrograde moving mitochondria of **B)** mean moving velocity (Axonal Location: p <0.0001; Directionality: p = 0.83; Interaction: p = 0.19) and **C)** relative frequency of time paused (Axonal Location: p = 0.0156; Directionality: p = 0.81; Interaction: p = 0.39). Means (solid line) ± SEM (shaded areas/error bars) were plotted for all graphs. Data were compared by two-way ANOVA and Šídák’s multiple comparisons tests. n=5 Mito.CFP mice.*p < 0.05, **p < 0.01, ***p < 0.001.

In summary, we have characterised a novel *in vivo* axonal transport profile of signalling endosomes and mitochondria across the NoR in multiple motor neuron sub-types within the sciatic nerve.

## Discussion

Axons are reliant upon efficient transport to maintain neuronal homeostasis. Using our intravital imaging approach (Sleigh et al., 2020b; Tosolini et al., 2021), we aimed to resolve the *in vivo* axonal transport dynamics of multiple organelles at the NoR in intact and synaptically-connected motor axons in the sciatic nerves of healthy mice. Firstly, we observed equivalent morphological features of the NoR in motor axons innervating prototypical fast and slow muscles. Next, we observed co-clustering of signalling endosomes and mitochondria specifically at distal side of the NoR. Our *in vivo* axonal transport analyses revealed a slowing of mitochondria and signalling endosomes as they approach the NoR, with a concomitant rise in pausing events. This was followed by an increase in velocity in the adjacent intranodal region, irrespective of directionality, organelle, and motor neuron subtype. Collectively, these findings further our understanding of the morphology and physiology of NoR in peripheral nerve axons.

### Peripheral nerve NoR morphology

We have previously reported that peripheral cholinergic motor axons transporting HCT upon uptake at neuromuscular junctions are larger in diameter than HCT-positive peripheral sensory axons innervating the same muscle (Sleigh et al., 2020a), and that ventral roots have a larger average calibre than dorsal roots in healthy mice (Rossor et al., 2020). Such differences have also been shown in central nervous system (CNS) axons (e.g. auditory system neurons; Ford et al., 2015) and in ventral funiculi axons in the thoracic spinal cord (Perrot et al., 2007). Conversely, the mean intranodal diameters of motor axons innervating the TA, lateral gastrocnemius and soleus muscles do not differ (Tosolini et al., 2022).

Here, we reveal comparable NoR morphological features between HCT-labelled FMN and SMN sciatic nerve axons, which are in turn, also comparable to the previously reported sciatic nerve profiles (Perrot et al., 2007) and morphometric results obtained from tibial nerve explants (Walker et al., 2019). In contrast, morphological features of the NoR in the CNS axons, such as those in the rat optic nerve and cerebral cortex, are heterogenous, with changes observed between axons rather than within axons, even when comparing axons with similar intranodal diameters (Arancibia-Cárcamo et al., 2017).

### Organelle accumulations at the NoR

We show for the first time dual-organelle accumulation at the NoR *in vivo* and from the same axon. As the peak fluorescence signal from both signalling endosomes and mitochondria was approximately 3-4.5 µm distally from the centre of the NoR, it is likely that these clusters were located in the distal juxtaparanode region. Indeed, the peak signal from the distal region was approximately 50% greater than the proximal internode for both organelles. Our data are in line with previous reports of expansions in the distal juxtaparanode of myelinated axons, which have been attributed to accumulations of diverse organelles, including retrogradely transported glycoproteins (Armstrong et al., 1987), retrogradely transported horseradish peroxidase (HRP) (Berthold and Mellström, 1982), multivesicular bodies (Berthold et al., 1993), lysosomes (Gatzinsky and Berthold, 1990) and mitochondria (Fabricius et al., 1993). However, the finding that mitochondria do not cluster at the NoR in small-diameter myelinated CNS axons (Edgar et al., 2008) and optic nerve (Perge et al., 2009), suggests there may be regional and/or neuronal subtype differences in organelle clusters at the NoR.

Mechanistically, it is unclear whether organelle accumulation at the NoR is an active process, or is simply a consequence of a localised cytoskeletal bottleneck. Although neurofilaments are reduced up to 10-fold at the NoR (Rydmark, 1981; Walker et al., 2019; Ciocanel et al., 2020), the microtubule network must be preserved to support axonal transport over the long distances covered by peripheral nerve axons (Hahn et al., 2019). Hence, the accumulation of organelles at the NoR might be structurally and functionally linked, with several potential mechanisms that could cause such focal accumulations, including differences in: 1) axonal protein translation (Rangaraju et al., 2017); 2) metabolic demands (Chiu, 2011); 3) axo-glial communication (Ronzano et al., 2021); 4) activity dependent alterations to microtubule structure (Peña-Ortega et al., 2022); 5) differential distributions of motor proteins and their regulators (Cason and Holzbaur, 2022); 6) hotspots for cytoskeletal anchoring (e.g., syntaphilin-mediated) (Kang et al., 2008); or 7) microtubule post-translational modifications (Janke and Magiera, 2020).

On the other hand, arrested mitochondrial motility is linked to activity-dependent axonal Ca^2+^ elevations (Zhang et al., 2010; Zhang and David, 2016). Furthermore, radial contractility alters axonal size to accommodate the movements of larger organelles (Pan et al., 2021). Thus, focal organelle accumulations might be linked to activity and can be transient in nature; however considerable follow up assessments are required to determine their mechanisms and dynamics. Moreover, direct interactions between Rab7-positive signalling endosomes and mitochondria, as well as co-transport with other organelles, could also influence nodal accumulation dynamics (Cioni et al., 2019; Liao et al., 2019; Obara et al., 2024).

### Axonal transport dynamics of diverse organelles at the NoR

While we have previously characterised signalling endosome and mitochondrial transport in physiological and pathological conditions (Bilsland et al., 2010; Sleigh et al., 2020a, 2024; Tosolini et al., 2022), this is the first study to quantitatively assess *in vivo* axonal transport dynamics of these organelles specifically at the NoR. The transport dynamics of signalling endosomes and mitochondria across the node were similar, with both organelles displaying faster velocities at the internode, and more pausing at the NoR, which contrasts with the report of transient accelerations of neurofilaments through nodal constrictions (Walker et al., 2019). We also identified faster speeds in the proximal internode compared to the distal internode of retrogradely transported signalling endosomes and anterogradely, but not retrogradely, moving mitochondria. Perhaps mechanistically linked, this transport phenotype is on the opposite internodal region to the organelle accumulations, which collectively suggests the presence of spatially-linked structural and functional differences at the NoR of peripheral nerves. Given the similar composition and function of the NoR and the axon initial segment (e.g., distribution of NaV1.6 channels (Boiko et al., 2003), Ank-G (Thome et al., 2023), and Neurofascin-186 (Hedstrom et al., 2007)), it would be interesting to compare transport dynamics between these two sites.

Both signalling endosomes and mitochondria, which undergo fast axonal transport, display a general slowing of transport speeds and an increase in pausing through the NoR, whereas neurofilaments, which undergo slow axonal transport, instead accelerate across the node (Walker et al., 2019). Such differences might be directly attributed to the machinery driving fast and slow axonal transport (Roy, 2020; Twelvetrees, 2020). However, despite both being classified as fast axonal transport, in adulthood signalling endosomes are transported faster than mitochondria *in vivo* (Bilsland et al., 2010; Sajic et al., 2013; Tosolini et al., 2022), which is likely linked to their divergent biological roles.

Signalling endosomes are retrogradely trafficked as Rab7-positive organelles, which function to propagate signalling from the periphery to the cell body to impact on translation and transcriptional events (Villarroel-Campos et al., 2018). This transport process can be modulated by several factors, including neurotrophins and their receptors (Deinhardt et al., 2006; Budzinska et al., 2020; Rhymes et al., 2022; Tosolini et al., 2022; Moya-Alvarado et al., 2023; Sleigh et al., 2023; Vargas et al., 2023; Rhymes et al., 2024) and components of the extracellular matrix (Surana et al., 2024). In contrast, mitochondrial transport ensures that specific axonal regions, such as synapses and NoRs, are enriched with sufficient mitochondria to respond to the localised high energy and Ca^2+^ handling demands (Sheng and Cai, 2012; Sun et al., 2013; Sheng, 2017).

In conclusion, using intravital imaging, we have characterised NoR morphologies in FMNs and SMNs, identified accumulations of signalling endosomes and mitochondria at the distal NoR, and determined the axonal transport dynamics of both organelles through the NoR. Finally, this work has clear implications for the peripheral nervous system and its numerous disorders (e.g., motor neuron diseases, multiple sclerosis) (Appeltshauser et al., 2023; Eshed-Eisenbach et al., 2023), and suggests that both the internodal and nodal transport modalities should be monitored in neuronal models of neurodegenerative diseases.

## Author Contributions

Conceptualisation: APT and GS. Investigation APT, FA, and SS, JNS. Writing and figure production: APT and GS, with input from all authors. Funding acquisition: APT and GS. All authors approved this submission.

## Declaration of Interests

The authors declare no competing interests.

## Acknowledgements

We thank the personnel of the Denny Brown Laboratories (Queen Square Institute of Neurology, University College London) for assistance in maintaining the mouse colonies, and Elena R. Rhymes and David Villarroel-Campos (Queen Square Institute of Neurology, University College London) for critical reading of the manuscript. This work was supported by a Junior Non-Clinical Fellowship from the Motor Neuron Disease Association (Tosolini/Oct20/973-799) (APT); a Col Bambrick MND Research Grant from Motor Neuron Disease Research Australia (IG 2450) (APT); a FightMND Drug Development Grant awarded to Giovanni Nardo (Istituto di Ricerche Farmacologiche Mario Negri IRCCS) (DDG-73; for APT); EMBO short-term fellowship (SN); Medical Research Council fellowships (MR/S006990/1 and MR/Y010949/1) (JNS); Wellcome Trust Senior Investigator Awards (107116/Z/15/Z and 223022/Z/21/Z) (GS), and a UK Dementia Research Institute award (UKDRI-1005) (GS).

## Notes

### Competing Interest Statement

The authors have declared no competing interest.

## References

Appeltshauser L, Linke J, Heil HS, Karus C, Schenk J, Hemmen K, Sommer C, Doppler K, Heinze KG (2023) Super-resolution imaging pinpoints the periodic ultrastructure at the human node of Ranvier and its disruption in patients with polyneuropathy. Neurobiol Dis 182:106139.

Arancibia-Cárcamo IL, Ford MC, Cossell L, Ishida K, Tohyama K, Attwell D (2017) Node of Ranvier length as a potential regulator of myelinated axon conduction speed Nave K-A, ed. eLife 6:e23329.

Armstrong R, Toews AD, Morell P (1987) Axonal transport through nodes of Ranvier. Brain Res 412:196–199.

Bercsenyi K, Schmieg N, Bryson JB, Wallace M, Caccin P, Golding M, Zanotti G, Greensmith L, Nischt R, Schiavo G (2014) Tetanus toxin entry. Nidogens are therapeutic targets for the prevention of tetanus. Science 346:1118–1123.

Berth SH, Lloyd TE (2023) Disruption of axonal transport in neurodegeneration. J Clin Invest 133.

Berthold C-H, Fabricius C, Rydmark M, Andersn B (1993) Axoplasmic organelles at nodes of Ranvier. I. Occurrence and distribution in large myelinated spinal root axons of the adult cat. J Neurocytol 22:925–940.

Berthold C-H, Mellström A (1982) Distribution of peroxidase activity at nodes of Ranvier after intramuscular administration of horseradish peroxidase in the cat. Neuroscience 7:45–54.

Bilsland LG, Sahai E, Kelly G, Golding M, Greensmith L, Schiavo G (2010) Deficits in axonal transport precede ALS symptoms in vivo. Proc Natl Acad Sci 107:20523–20528.

Blum JA, Klemm S, Shadrach JL, Guttenplan KA, Nakayama L, Kathiria A, Hoang PT, Gautier O, Kaltschmidt JA, Greenleaf WJ, Gitler AD (2021) Single-cell transcriptomic analysis of the adult mouse spinal cord reveals molecular diversity of autonomic and skeletal motor neurons. Nat Neurosci 24:572–583.

Boiko T, Van Wart A, Caldwell JH, Levinson SR, Trimmer JS, Matthews G (2003) Functional specialization of the axon initial segment by isoform-specific sodium channel targeting. J Neurosci Off J Soc Neurosci 23:2306–2313.

Brown A (2003) Axonal transport of membranous and nonmembranous cargoes. J Cell Biol 160:817–821.

Budzinska MI, Campos DV, Golding M, Weston A, Collinson L, Snijders AP, Schiavo G (2020) PTPN23 binds the dynein adaptor BICD1 and is required for endocytic sorting of neurotrophin receptors. J Cell Sci 133:jcs242412–14.

Cason SE, Holzbaur ELF (2022) Selective motor activation in organelle transport along axons. Nat Rev Mol Cell Biol:1–16.

Chiu SY (2011) Matching Mitochondria to Metabolic Needs at Nodes of Ranvier. The Neuroscientist 17:343–350.

Ciocanel M-V, Jung P, Brown A (2020) A mechanism for neurofilament transport acceleration through nodes of Ranvier. Mol Biol Cell 31:640–654.

Cioni J-M, Lin JQ, Holtermann AV, Koppers M, Jakobs MAH, Azizi A, Turner-Bridger B, Shigeoka T, Franze K, Harris WA, Holt CE (2019) Late Endosomes Act as mRNA Translation Platforms and Sustain Mitochondria in Axons. Cell 176:56–72.e15.

Deinhardt K, Salinas S, Verastegui C, Watson R, Worth D, Hanrahan S, Bucci C, Schiavo G (2006) Rab5 and Rab7 Control Endocytic Sorting along the Axonal Retrograde Transport Pathway. Neuron 52:293–305.

D’Este E, Kamin D, Balzarotti F, Hell SW (2017) Ultrastructural anatomy of nodes of Ranvier in the peripheral nervous system as revealed by STED microscopy. Proc Natl Acad Sci 114:E191–E199.

Edgar JM, McCulloch MC, Thomson CE, Griffiths IR (2008) Distribution of mitochondria along small-diameter myelinated central nervous system axons. J Neurosci Res 86:2250–2257.

Ershov D, Phan M-S, Pylvänäinen JW, Rigaud SU, Blanc LL, Charles-Orszag A, Conway JRW, Laine RF, Roy NH, Bonazzi D, Duménil G, Jacquemet G, Tinevez J-Y (2022) TrackMate 7: integrating state-of-the-art segmentation algorithms into tracking pipelines. Nat Methods 19:829–832.

Eshed-Eisenbach Y, Brophy PJ, Peles E (2023) Nodes of Ranvier in health and disease. J Peripher Nerv Syst JPNS 28 Suppl 3:S3–S11.

Fabricius C, Berthold C-H, Rydmark M (1993) Axoplasmic organelles at nodes of Ranvier. II. Occurrence and distribution in large myelinated spinal cord axons of the adult cat. J Neurocytol 22:941–954.

Ford MC, Alexandrova O, Cossell L, Stange-Marten A, Sinclair J, Kopp-Scheinpflug C, Pecka M, Attwell D, Grothe B (2015) Tuning of Ranvier node and internode properties in myelinated axons to adjust action potential timing. Nat Commun 6:8073.

Gatzinsky KP, Berthold C-H (1990) Lysosomal activity at nodes of Ranvier during retrograde axonal transport of horseradish peroxidase in alpha-motor neurons of the cat. J Neurocytol 19:989–1002.

Gibbs KL, Greensmith L, Schiavo G (2015) Regulation of Axonal Transport by Protein Kinases. Trends Biochem Sci 40:597–610.

Gibbs KL, Kalmar B, Sleigh JN, Greensmith L, Schiavo G (2016) In vivo imaging of axonal transport in murine motor and sensory neurons. J Neurosci Methods 257:26–33.

Gong S, Zheng C, Doughty ML, Losos K, Didkovsky N, Schambra UB, Nowak NJ, Joyner A, Leblanc G, Hatten ME, Heintz N (2003) A gene expression atlas of the central nervous system based on bacterial artificial chromosomes. Nature 425:917–925.

Goto-Silva L, McShane MP, Salinas S, Kalaidzidis Y, Schiavo G, Zerial M (2019) Retrograde transport of Akt by a neuronal Rab5-APPL1 endosome. Sci Rep 9:2433.

Hahn I, Voelzmann A, Liew Y-T, Costa-Gomes B, Prokop A (2019) The model of local axon homeostasis - explaining the role and regulation of microtubule bundles in axon maintenance and pathology. Neural Develop:1–28.

Hedstrom KL, Xu X, Ogawa Y, Frischknecht R, Seidenbecher CI, Shrager P, Rasband MN (2007) Neurofascin assembles a specialized extracellular matrix at the axon initial segment. J Cell Biol 178:875–886.

Hsieh S, Kidd G, Crawford T, Xu Z, Lin W, Trapp B, Cleveland D, Griffin J (1994) Regional modulation of neurofilament organization by myelination in normal axons. J Neurosci 14:6392–6401.

Janke C, Magiera MM (2020) The tubulin code and its role in controlling microtubule properties and functions. Nat Rev Mol Cell Biol 21:307–326.

Johnson C, Holmes WR, Brown A, Jung P (2015) Minimizing the caliber of myelinated axons by means of nodal constrictions. J Neurophysiol 114:1874–1884.

Kalinski AL et al. (2019) Deacetylation of Miro1 by HDAC6 blocks mitochondrial transport and mediates axon growth inhibition. J Cell Biol 218:1871–1890.

Kang J-S, Tian J-H, Pan P-Y, Zald P, Li C, Deng C, Sheng Z-H (2008) Docking of Axonal Mitochondria by Syntaphilin Controls their Mobility and Affects Short-term Facilitation. Cell 132:137–148.

Lang Q, Schiavo G, Sleigh JN (2023) In vivo imaging of axonal transport in peripheral nerves of rodent forelimbs. Neuronal Signal 7:NS20220098.

Liao Y-C et al. (2019) RNA Granules Hitchhike on Lysosomes for Long-Distance Transport, Using Annexin A11 as a Molecular Tether. Cell 179:147–164.e20.

Maday S, Twelvetrees AE, Moughamian AJ, Holzbaur ELF (2014) Axonal Transport: Cargo-Specific Mechanisms of Motility and Regulation. Neuron 84:292–309.

Mallon BS, Shick HE, Kidd GJ, Macklin WB (2002) Proteolipid promoter activity distinguishes two populations of NG2-positive cells throughout neonatal cortical development. J Neurosci Off J Soc Neurosci 22:876–885.

Misgeld T, Kerschensteiner M, Bareyre FM, Burgess RW, Lichtman JW (2007) Imaging axonal transport of mitochondria in vivo. Nat Methods 4:559–561.

Mohan R, Tosolini AP, Morris R (2014) Targeting the motor end plates in the mouse hindlimb gives access to a greater number of spinal cord motor neurons: An approach to maximize retrograde transport. Neuroscience 274:318–330.

Mohan R, Tosolini AP, Morris R (2015) Intramuscular Injections Along the Motor End Plates: A Minimally Invasive Approach to Shuttle Tracers Directly into Motor Neurons. J Vis Exp JoVE:1–8.

Moya-Alvarado G, Tiburcio-Felix R, Ibáñez MR, Aguirre-Soto AA, Guerra MV, Wu C, Mobley WC, Perlson E, Bronfman FC (2023) BDNF/TrkB signaling endosomes in axons coordinate CREB/mTOR activation and protein synthesis in the cell body to induce dendritic growth in cortical neurons. eLife 12:e77455.

Negro S, Lauria F, Stazi M, Tebaldi T, D’Este G, Pirazzini M, Megighian A, Lessi F, Mazzanti CM, Sales G, Romualdi C, Fillo S, Lista F, Sleigh JN, Tosolini AP, Schiavo G, Viero G, Rigoni M (2022) Hydrogen peroxide induced by nerve injury promotes axon regeneration via connective tissue growth factor. Acta Neuropathol Commun 10:189.

Negro S, Lessi F, Duregotti E, Aretini P, Ferla ML, Franceschi S, Menicagli M, Bergamin E, Radice E, Thelen M, Megighian A, Pirazzini M, Mazzanti CM, Rigoni M, Montecucco C (2017) CXCL12α/SDF‐1 from perisynaptic Schwann cells promotes regeneration of injured motor axon terminals. EMBO Mol Med 9:1000–1010.

Nirschl JJ, Ghiretti AE, Holzbaur ELF (2017) The impact of cytoskeletal organization on the local regulation of neuronal transport. Nat Publ Group 18:585–597.

Obara CJ, Nixon-Abell J, Moore AS, Riccio F, Hoffman DP, Shtengel G, Xu CS, Schaefer K, Pasolli HA, Masson J-B, Hess HF, Calderon CP, Blackstone C, Lippincott-Schwartz J (2024) Motion of VAPB molecules reveals ER–mitochondria contact site subdomains. Nature 626:169–176.

Pan X, Zhou Y, Hotulainen P, Meunier FA, Wang T (2021) The axonal radial contractility: Structural basis underlying a new form of neural plasticity. BioEssays 43:2100033.

Peña-Ortega F, Robles-Gómez ÁA, Xolalpa-Cueva L (2022) Microtubules as Regulators of Neural Network Shape and Function: Focus on Excitability, Plasticity and Memory. Cells 11:923.

Perge JA, Koch K, Miller R, Sterling P, Balasubramanian V (2009) How the optic nerve allocates space, energy capacity, and information. J Neurosci Off J Soc Neurosci 29:7917–7928.

Perrot R, Lonchampt P, Peterson AC, Eyer J (2007) Axonal neurofilaments control multiple fiber properties but do not influence structure or spacing of nodes of Ranvier. J Neurosci Off J Soc Neurosci 27:9573–9584.

Ragagnin AMG, Shadfar S, Vidal M, Jamali MS, Atkin JD (2019) Motor Neuron Susceptibility in ALS/FTD. Front Neurosci 13:532.

Rangaraju V, Dieck S tom, Schuman EM (2017) Local translation in neuronal compartments: how local is local? EMBO Rep 18:693–711.

Rasband MN, Peles E (2021) Mechanisms of node of Ranvier assembly. Nat Rev Neurosci 22:7–20.

Reles A, Friede RL (1991) Axonal cytoskeleton at the nodes of Ranvier. J Neurocytol 20:450–458.

Restani L, Giribaldi F, Manich M, Bercsenyi K, Menendez G, Rossetto O, Caleo M, Schiavo G (2012) Botulinum Neurotoxins A and E Undergo Retrograde Axonal Transport in Primary Motor Neurons Blanke SR, ed. PLoS Pathog 8:e1003087.

Rhymes ER, Simkin RL, Qu J, Villarroel-Campos D, Surana S, Tong Y, Shapiro R, Burgess RW, Yang X-L, Schiavo G, Sleigh JN (2024) Boosting BDNF in muscle rescues impaired axonal transport in a mouse model of DI-CMTC peripheral neuropathy. Neurobiol Dis 195:106501.

Rhymes ER, Tosolini AP, Fellows AD, Mahy W, McDonald NQ, Schiavo G (2022) Bimodal regulation of axonal transport by the GDNF-RET signalling axis in healthy and diseased motor neurons. Cell Death Dis 13:584.

Rodella U, Negro S, Scorzeto M, Bergamin E, Jalink K, Montecucco C, Yuki N, Rigoni M (2017) Schwann cells are activated by ATP released from neurons in an in vitro cellular model of Miller Fisher syndrome. Dis Model Mech 10:597–603.

Ronzano R, Roux T, Thetiot M, Aigrot MS, Richard L, Lejeune FX, Mazuir E, Vallat JM, Lubetzki C, Desmazières A (2021) Microglia-neuron interaction at nodes of Ranvier depends on neuronal activity through potassium release and contributes to remyelination. Nat Commun 12:5219.

Rossor AM, Sleigh JN, Groves M, Muntoni F, Reilly MM, Hoogenraad CC, Schiavo G (2020) Loss of BICD2 in muscle drives motor neuron loss in a developmental form of spinal muscular atrophy. Acta Neuropathol Commun 8:34–12.

Roy S (2020) Finding order in slow axonal transport. Curr Opin Neurobiol 63:87–94.

Rydmark M (1981) Nodal axon diameter correlates linearly with internodal axon diameter in spinal roots of the cat. Neurosci Lett 24:247–250.

Sajic M, Mastrolia V, Lee CY, Trigo D, Sadeghian M, Mosley AJ, Gregson NA, Duchen MR, Smith KJ (2013) Impulse Conduction Increases Mitochondrial Transport in Adult Mammalian Peripheral Nerves In Vivo. PLOS Biol 11:e1001754.

Sheng Z-H (2017) The Interplay of Axonal Energy Homeostasis and Mitochondrial Trafficking and Anchoring. Trends Cell Biol 27:403–416.

Sheng Z-H, Cai Q (2012) Mitochondrial transport in neurons: impact on synaptic homeostasis and neurodegeneration. Nat Rev Neurosci 13:77–93.

Sleigh J, Mattedi F, Richter S, Annuario E, Ng K, Steinmark IE, Ivanova I, Darabán I, Joshi P, Rhymes E, Awale S, Yahioglu G, Mitchell J, Suhling K, Schiavo G, Vagnoni A (2024) Age-specific and compartment-dependent changes in mitochondrial homeostasis and cytoplasmic viscosity in mouse peripheral neurons. Aging Cell [in press].

Sleigh JN, Rossor AM, Fellows AD, Tosolini AP, Schiavo G (2019) Axonal transport and neurological disease. Nat Rev Neurol 15:691–703.

Sleigh JN, Tosolini AP, Gordon D, Devoy A, Fratta P, Fisher EMC, Talbot K, Schiavo G (2020a) Mice Carrying ALS Mutant TDP-43, but Not Mutant FUS, Display In Vivo Defects in Axonal Transport of Signaling Endosomes. Cell Rep 30:3655–3662.e2.

Sleigh JN, Tosolini AP, Schiavo G (2020b) In Vivo Imaging of Anterograde and Retrograde Axonal Transport in Rodent Peripheral Nerves. Methods Mol Biol Clifton NJ 2143:271–292.

Sleigh JN, Villarroel-Campos D, Surana S, Wickenden T, Tong Y, Simkin RL, Vargas JNS, Rhymes ER, Tosolini AP, West SJ, Zhang Q, Yang X-L, Schiavo G (2023) Boosting peripheral BDNF rescues impaired in vivo axonal transport in CMT2D mice. JCI Insight.

Stifani N (2014) Motor neurons and the generation of spinal motor neuron diversity. Front Cell Neurosci 8:293.

Sun T, Qiao H, Pan P-Y, Chen Y, Sheng Z-H (2013) Motile Axonal Mitochondria Contribute to the Variability of Presynaptic Strength. Cell Rep 4:413–419.

Surana S, Tosolini AP, Meyer IFG, Fellows AD, Novoselov SS, Schiavo G (2018) The travel diaries of tetanus and botulinum neurotoxins. Toxicon 147:58–67.

Surana S, Villarroel-Campos D, Rhymes E, Kalyukina M, Panzi C, Novoselov S, Fabris F, Richter S, Pirazzini M, Zanotti G, Sleigh J, Schiavo G (2024) The tyrosine phosphatases LAR and PTPRD act as receptors of the nidogen-tetanus toxin complex EMBO J (In press).

Thome C, Janssen JM, Karabulut S, Acuna C, D’Este E, Soyka SJ, Baum K, Bock M, Lehmann N, Hasegawa M, Ganea DA, Benoit CM, Gründemann J, Schultz C, Bennett V, Jenkins PM, Engelhardt M (2023) Live imaging of excitable axonal microdomains in ankyrin-G-GFP mice. eLife 12.

Tosolini AP, Sleigh JN, Surana S, Rhymes ER, Cahalan SD, Schiavo G (2022) BDNF-dependent modulation of axonal transport is selectively impaired in ALS. Acta Neuropathol Commun 10:121.

Tosolini AP, Villarroel-Campos D, Schiavo G, Sleigh JN (2021) Expanding the Toolkit for In Vivo Imaging of Axonal Transport. J Vis Exp:e63471.

Tsukita S, Ishikawa H (1981) The cytoskeleton in myelinated axons: serial section study. Biomed Res 2:424–437.

Twelvetrees AE (2020) The lifecycle of the neuronal microtubule transport machinery. Semin Cell Dev Biol 107:74–81.

Vargas JNS et al. (2023) BDNF controls phosphorylation and transcriptional networks governing cytoskeleton organization and axonal regeneration. BioRxiv.

Vargas JNS, Sleigh JN, Schiavo G (2022) Coupling axonal mRNA transport and local translation to organelle maintenance and function. Curr Opin Cell Biol 74:97–103.

Villarroel-Campos D, Schiavo G, Lazo OM (2018) The many disguises of the signalling endosome. FEBS Lett 592:3615–3632.

Walker CL, Uchida A, Li Y, Trivedi N, Fenn JD, Monsma PC, Lariviere RC, Julien J-P, Jung P, Brown A (2019) Local Acceleration of Neurofilament Transport at Nodes of Ranvier. J Neurosci Off J Soc Neurosci 39:663–677.

Wang T, Li W, Martin S, Papadopulos A, Joensuu M, Liu C, Jiang A, Shamsollahi G, Amor R, Lanoue V, Padmanabhan P, Meunier FA (2020) Radial contractility of actomyosin rings facilitates axonal trafficking and structural stability. J Cell Biol 219:e201902001.

Zhang CL, Ho PL, Kintner DB, Sun D, Chiu SY (2010) Activity-Dependent Regulation of Mitochondrial Motility by Calcium and Na/K-ATPase at Nodes of Ranvier of Myelinated Nerves. J Neurosci 30:3555–3566.

Zhang Z, David G (2016) Stimulation‐induced Ca2+ influx at nodes of Ranvier in mouse peripheral motor axons. J Physiol 594:39–57.

