## Supplementary Material for "The node of Ranvier influences the *in vivo* axonal transport of mitochondria and signalling endosomes"

### Extended Data - Figures and Legends

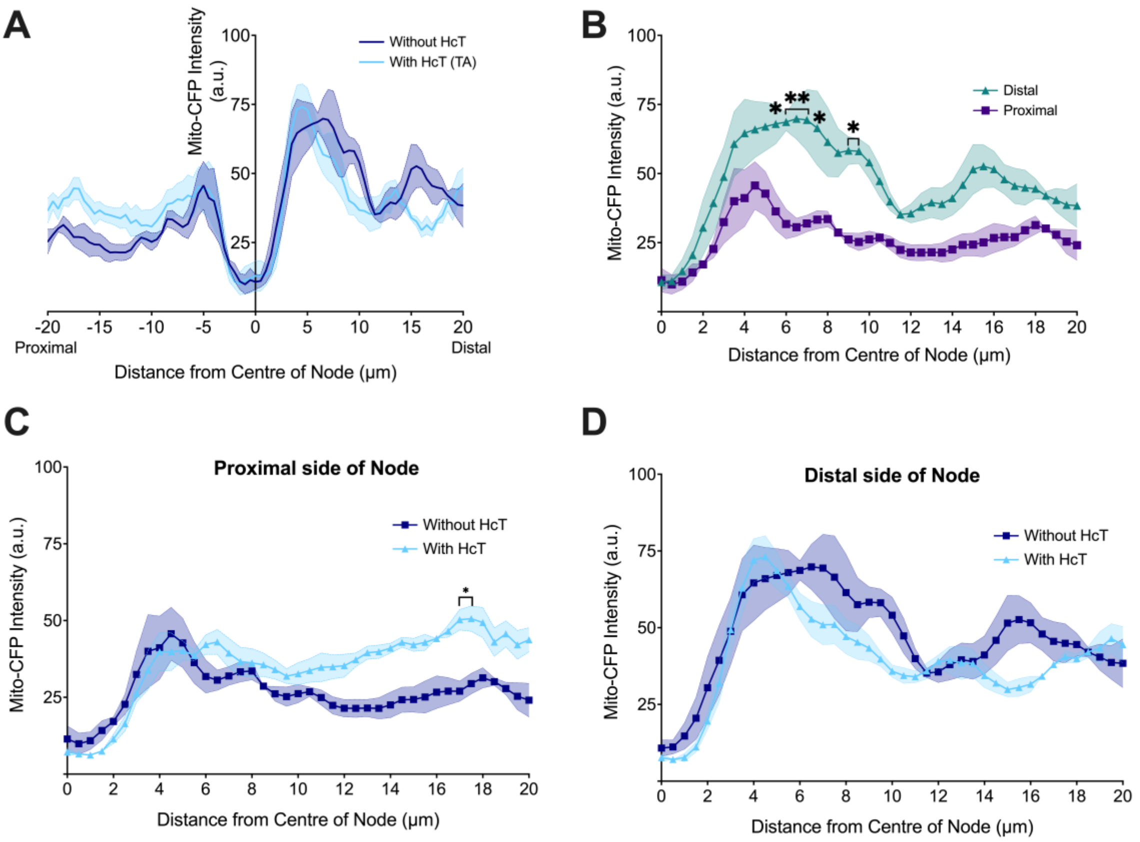


**Supplementary Figure 1. Mitochondrial clustering distal to the node of Ranvier is independent of $H_CT$ administration. A)** In sciatic nerve axons, we observed similar patterns of fluorescence intensity and localisation of Mito.CFP labelled mitochondria in experiments with (cyan) or without (navy) intramuscular $H_CT$ injections. **B)** In mice without intramuscular injections, we observed similar increases in mean fluorescence intensities of Mito.CFP labelled mitochondria on the distal side of the NoR. Comparing 'with $H_CT$' (cyan) vs 'without $H_CT$' (navy) indicates similar Mito.CFP mean fluorescence patterns on both the **C)** proximal (Axonal Location: $p < 0.0001$; Mean Relative Fluorescence: $p < 0.0001$; Interaction: $p < 0.0001$); and **D)** distal (Axonal Location: $p < 0.0001$; Mean Relative Fluorescence: $p < 0.0001$; Interaction: $p = 0.7746$) sides of the NoR. Means (solid line) ± SEM (shaded areas) were plotted for all graphs, n=5 animals, 16 axons. Data were compared by two-way ANOVA and Šídák's multiple comparisons tests. *$p < 0.05$. **Linked to Figure 2**.

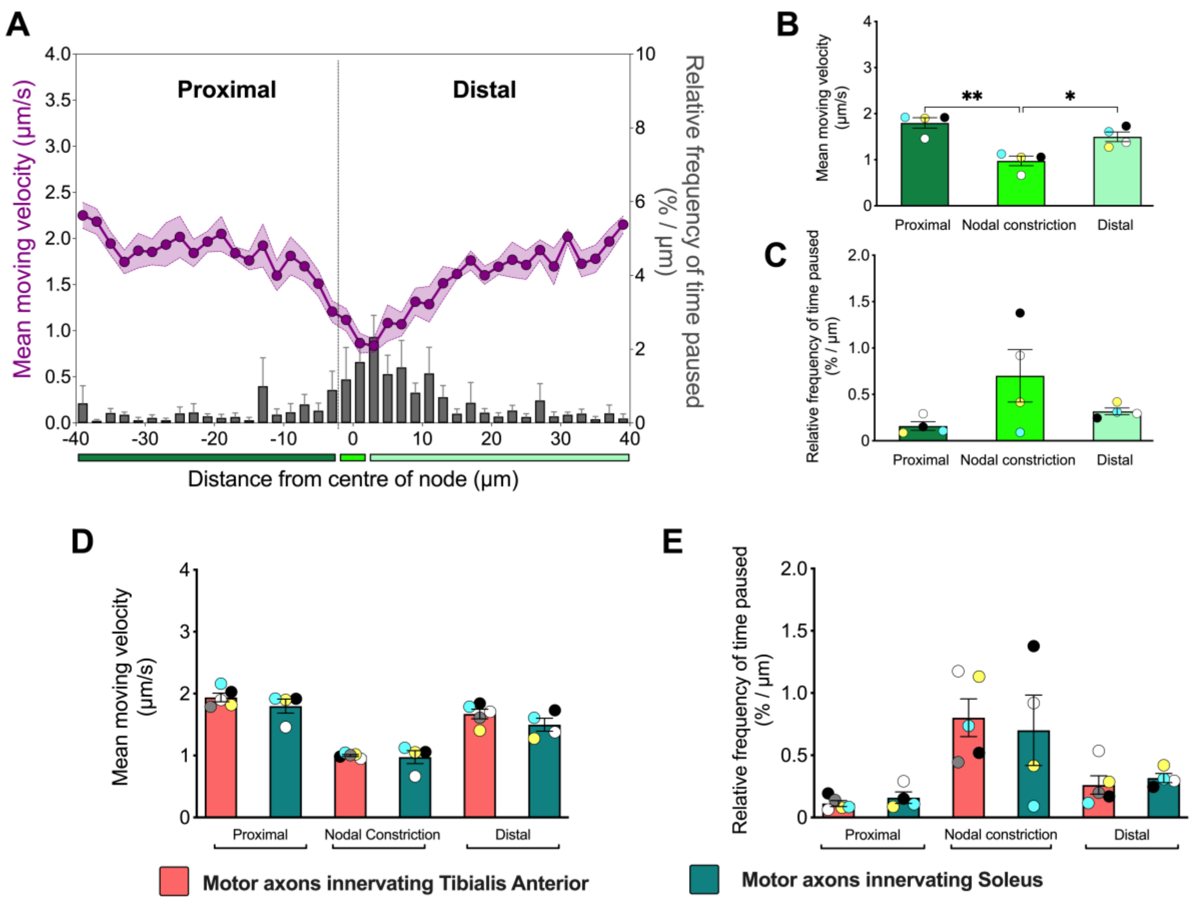


**Supplementary Figure 2. Signalling endosome dynamics through the node are similar between TA- and soleus-innervating motor axons – Linked to Figure 5. A)** Retrograde axonal transport dynamics (mean moving velocity [purple] and relative frequency of mean pausing [grey bars]) of $H_CT$-containing signalling endosomes across the nodal constriction (80 µm distance) in motor axons innervating the soleus muscle. The x-axis represents the distance from the centre of the node of Ranvier (µm) and is split into three segments: 1) *Proximal* = 38 µm of the proximal internode (dark green), 2) *Centre* = 4 µm representing the mean nodal length, as determined in **Figure 1D** (bright green), and 3) *Distal* = 38 µm of the distal internode (light green). Comparisons across the proximal internode (dark green), nodal constriction (bright green) and distal internode (light green) of the **B)** mean moving velocity ($p = 0.0014$), and **C)** relative frequency of mean pausing ($p = 0.11$). Comparisons from motor axons innervating the tibialis anterior (salmon) and soleus (teal) of the **D)** mean moving velocity ($p < 0.0001$) and **E)** relative frequency of mean pausing ($p = 0.0031$). Data were compared by an ordinary one-way ANOVA, followed by Holm-Šídák's multiple comparisons test. Means (solid line) ± SEM (shaded area/error bars) were plotted for all graphs. n=4 (soleus), n=5 (tibialis anterior) from Mito.CFP mice.*$p < 0.05$, **$p < 0.01$.

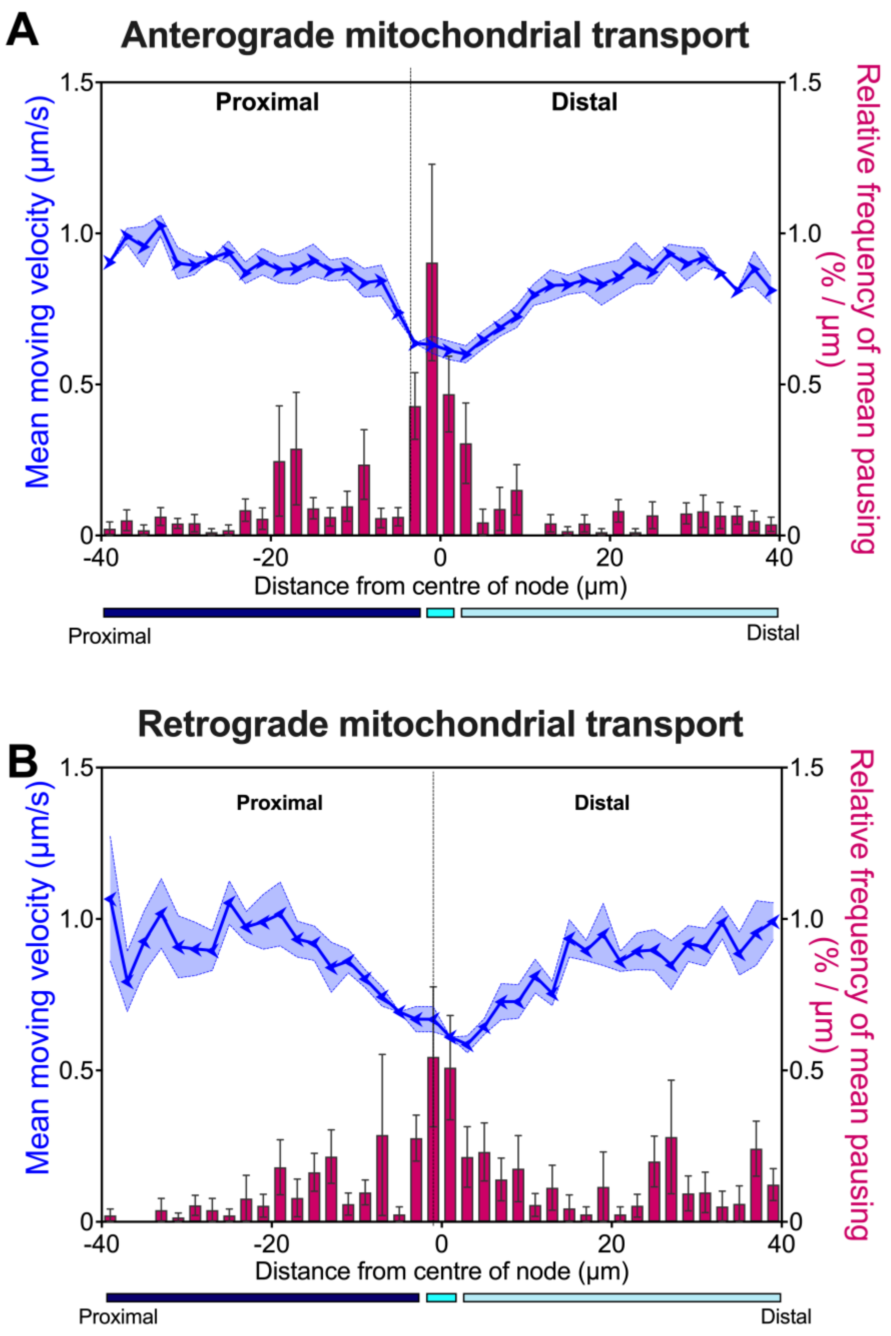


**Supplementary Figure 3. Mitochondria slow and pause more at the nodal constriction in both anterograde and retrograde directions – Linked to Figure 6.** *In vivo* mitochondrial **A)** anterograde and **B)** retrograde transport dynamics (mean moving velocity [blue arrowheads] and relative frequency of mean pausing [pink bars]) across the nodal constriction (80 µm distance) in axons innervating the tibialis anterior muscle. The x-axis represents the distance from the centre of the node of Ranvier (µm) and is split into three segments: 1) *Proximal* = 38 µm of the proximal internode (navy); 2) *Centre* = 4 µm representing the mean nodal length (as determined in **Figure 1D**; cyan); and 3) *Distal* = 38 µm of the distal internode (light blue). Means (solid line) ± SEM (shaded areas/error bars) were plotted for all graphs. n=5 from Mito.CFP mice.

**Extended Data – Movie Legends**

**Movie 1. $Na_V$1.6 clusters are located in the middle of the nodal constriction and are flanked by CAPSR-positive paranodal regions in a peripheral nerve cholinergic motor axon.** Representative videos of **A)** a rotating max projection z-stack, and **B)** slice-by-slice through the z-axis, highlighting the *i)* cholinergic motor axon, the immunolabelled *ii)* nodal ($Na_V$1.6) and *iii)* paranodal (CASPR) regions, along with the *iv)* overlay, from teased axons from a ChAT.eGFP mice sciatic nerve. Scale bar = 5 µm.

**Movie 2. *In vivo* axonal transport of mitochondria and signalling endosomes is observed through the nodal constrictions of a single sciatic nerve axon.** Representative time-lapse microscopy video of dual-organelle *in vivo* axonal transport through the nodal constriction from a Mito.CFP mouse. Frame interval = 1.44 s; number of frames: 201; Playback rate: 15 fps; scale bar = 10 µm.

**Movie 3. Signalling endosomes that traverse the nodal constrictions are not present in the Schwann cell cytosol.** Representative video of a **A)** rotating max projection z-stack, and **B)** slice-by-slice through the z-axis, highlighting the *i)* $H_C$T-containing signalling endosomes (magenta), within the *ii)* Schwann cell cytosol (eGFP), and *iii)* overlay, from the same teased axon of a PLP-eGFP mice sciatic nerve. Scale bar = 10 µm.

**Movie 4. Representative video of *in vivo* axonal transport of HcT-containing signalling endosomes traversing the node of Ranvier in a single sciatic motor axon.** Top panel = ChAT.eGFP signal; Middle panel = retrogradely transporting HcT-containing signalling endosomes; Bottom panel = overlay. Frame interval: 0.53 s; number of frames: 400; playback rate: 20 fps; scale bar = 10 µm.

**Movie 5. Retrograde axonal transport of HcT (magenta) can display an unusual trajectory through the node of Ranvier in peripheral nerve motor axons (green) – Linked to Figure 4A.** Frame interval: 0.45 s; playback rate = 25 fps; acquisition time = 41 s; and scale bar: 10 µm

**Movie 6. Retrograde axonal transport of HcT-containing signalling endosomes (magenta) travelling in a circular path (green arrow) around mitochondria (cyan) on the distal side of the node of Ranvier – Linked to Figure 4B.** Frame interval: 1.64 s; playback rate = 8 fps; acquisition time = 3 mins 11 seconds; scale bar = 10 µm

**Movie 7. Representative videos of tracked mitochondria separated by directionality – Linked to Figure 6 and Supplementary Figure 3. A)** Representative unprocessed time-lapse video. Using the TrackMate plugin in FIJI/ImageJ, mitochondria (green dots and lines) were manually tracked in the **B)** anterograde (magenta) and **C)** retrograde (gold) directions. Frame interval: 1.77 s; playback rate = 20 fps; acquisition time = 10 m 21 s; scale bar: 10 µm.
